# Mapping protein interactions of sodium channel Na_V_1.7 using epitope-tagged gene targeted mice

**DOI:** 10.1101/118497

**Authors:** Alexandros H. Kanellopoulos, Jennifer Koenig, Honglei Huang, Martina Pyrski, Queensta Millet, Stephane Lolignier, Toru Morohashi, Samuel J. Gossage, Maude Jay, John Linley, Georgios Baskozos, Benedikt Kessler, James J. Cox, Frank Zufall, John N. Wood, Jing Zhao

## Abstract

The voltage-gated sodium channel Na_V_1.7 plays a critical role in pain pathways. Besides action potential propagation, Na_V_1.7 regulates neurotransmitter release, integrates depolarizing inputs over long periods and regulates transcription. In order to better understand these functions, we generated an epitope-tagged Na_V_1.7 mouse that showed normal pain behavior. Analysis of Na_V_1.7 complexes affinity-purified under native conditions by mass spectrometry revealed 267 Na_V_1.7 associated proteins including known interactors, such as the sodium channel β3 subunit (Scn3b) and collapsin response mediator protein (Crmp2), and novel interactors. Selected novel Na_V_1.7 protein interactors membrane-trafficking protein synapototagmin-2 (Syt2), G protein-regulated inducer of neurite outgrowth 1 (Gprin1), L-type amino acid transporter 1 (Lat1) and transmembrane P24 trafficking protein 10 (Tmed10) together with Scn3b and Crmp2 were validated using co-immunoprecipitation and functional assays. The information provided with this physiologically normal epitope-tagged mouse should provide useful insights into the pain mechanisms associated with Na_V_1.7 channel function.

## Introduction

Pain is a major clinical problem. A 2012 National Health Interview Survey (NHIS) in the United States revealed that 25.3 million adults (11.2%) suffered from daily (chronic) pain and 23.4 million (10.3%) reported severe pain within a previous three-month period (Nahin, 2015). Many types of chronic pain are difficult to treat as most available drugs have limited efficacy and can cause side effects. There is thus a huge unmet need for novel analgesics (Woodcock, 2009). Recent human and animal genetic studies have indicated that the voltage-gated sodium channel (VGSC) Na_V_1.7 plays a crucial role in pain signaling (Dib-Hajj et al., 2013, Cox et al., 2006, Nassar et al., 2004), highlighting Na_V_1.7 as a promising drug target for development of novel analgesics (Emery et al., 2016, Habib et al., 2015).

VGSCs consist of a large pore-forming α-subunit (~260 kDa) together with associated auxiliary β-subunits (33-36 kDa) and play a fundamental role in the initiation and propagation of action potentials in electrically excitable cells. Nine isoforms of α-subunits (Na_V_1.1-1.9) that display distinct expression patterns and variable channel properties have been identified in mammals (Frank and Catterall, 2003). Na_V_1.7, encoded by the gene *SCN9A* in humans, is selectively expressed peripherally in dorsal root ganglion (DRG), trigeminal ganglia and sympathetic neurons (Black et al., 2012, Toledo-Aral et al., 1997), as well as the central nervous system (Branco et al., 2016, Weiss et al., 2011). As a large membrane ion channel, Na_V_1.7 produces a fast-activating, fast-inactivating and slowly re-priming current (Klugbauer et al., 1995), acting as a threshold channel to contribute to the generation and propagation of action potentials by amplifying small sub-threshold depolarisations (Rush et al., 2007). The particular electrophysiological characteristics of Na_V_1.7 suggests that it plays a key role in initiating action potentials in response to depolarization of sensory neurons by noxious stimuli (Habib et al., 2015). In animal studies, our previous results demonstrated that conditional Na_V_1.7 knockout mice have major deficits in acute, inflammatory and neuropathic pain (Minett et al., 2012, Nassar et al., 2004). Human genetic studies showed that loss of function mutations in Na_V_1.7 lead to congenital insensitivity to pain whereas gain of function mutations cause a range of painful inherited disorders (Dib-Hajj et al., 2013, Cox et al., 2006). Patients with recessive Na_V_1.7 mutations are normal (apart from being pain-free and anosmic), suggesting selective Na_V_1.7 blocking drugs are therefore likely to be effective analgesics with limited side effects (Cox et al., 2006). Recent studies also showed that Na_V_1.7 is involved in neurotransmitter release in both the olfactory bulb and spinal cord (Minett et al., 2012, Weiss et al., 2011).

Over the past decade, an enormous effort has been made to develop selective Na_V_1.7 blockers. Efficient selective Na_V_1.7 antagonists have been developed, however, Na_V_1.7 antagonists require co-administration of opioids to be fully analgesic (Minett et al., 2015). Na_V_1.7 thus remains an important analgesic drug target and an alternative strategy to target Na_V_1.7 could be to interfere with either intracellular trafficking of the channel to the plasma membrane, or the downstream effects of the channel on opioid peptide expression. Although some molecules, such as Crmp2, ubiquitin ligase Nedd4-2, fibroblast growth factor 13 (FGF13) and PDZ domain-containing protein Pdzd2, have been associated with the regulation of trafficking and degradation of Na_V_1.7, the entire protein-protein interaction network of Na_V_1.7 still needs to be defined (Dustrude et al., 2013, Laedermann et al., 2013, Shao et al., 2009, Bao, 2015, Yang et al., 2017).Affinity purification (AP) and targeted tandem affinity purification (TAP) combined with mass spectrometry (MS) is a useful method for mapping the organization of multiprotein complexes (Wildburger et al., 2015, Angrand et al., 2006). Using the AP-MS or TAP-MS method to characterize protein complexes from transgenic mice allows the identification of complexes in their native physiological environment in contact with proteins that might only be specifically expressed in certain tissues (Fernandez et al., 2009). The principal aim of this study was to identify new protein interaction partners and networks involved in the trafficking and regulation of sodium channel Na_V_1.7 using mass spectrometry and our recently generated epitope (TAP)-tagged Na_V_1.7 knock-in mouse. Such information should throw new light on the mechanisms through which Na_V_1.7 regulates a variety of physiological systems, as well as propagating action potentials, especially in pain signaling pathways.

## Results

### The TAP tag does not affect Na_V_1.7 channel function

Prior to generating the TAP-tagged Na_V_1.7 knock-in mouse, we tested the channel function of TAP-tagged Na_V_1.7 in HEK293 cells by establishing a HEK293 cell line stably expressing TAP-tagged Na_V_1.7. A 5-kDa TAP tag consisting of a polyhistidine affinity tag (HAT) and a 3x FLAG tag in tandem (Terpe, 2003), separated by a unique TEV-protease cleavage site, was fused to the C-terminus of Na_V_1.7 (Fig 1A). The expression of TAP-tagged Na_V_1.7 was detected with immunocytochemistry using an anti-FLAG antibody. The result showed that all the HEK293 cells expressed TAP-tagged Na_V_1.7 (Fig 1B). The channel function of TAP-tagged Na_V_1.7 was examined with electrophysiological analysis and the result showed that all the cells presented normal functional Na_V_1.7 currents (Fig 1C). Activation and fast inactivation data were identical for wild-type Na_V_1.7 and TAP-tagged Na_V_1.7 (Fig 1D) demonstrating that the TAP tag does not affect channel function of Na_V_1.7.

**Figure 1.**
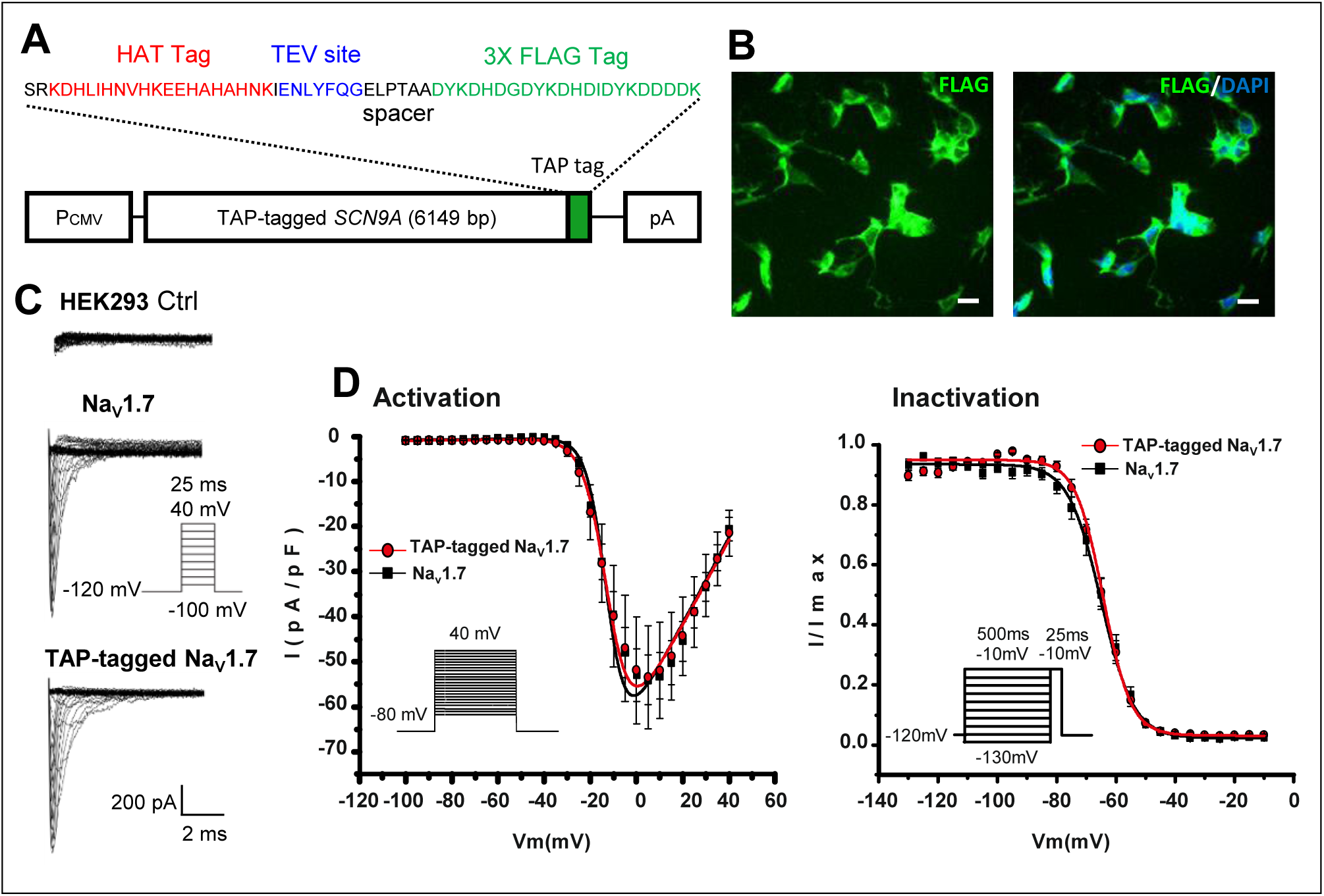
Characterisation of TAP-tagged Na_V_1.7 in HEK293 cells. A The diagram of TAP-tagged *SCN9A* cDNA construct used to establish the stably expressing TAP-tagged Na_v_1.7 HEK293 cell line. A sequence encoding a TAP tag was cloned immediately upstream of the stop codon of *SCN9A* coding the wild-type Na_v_1.7. B Left panel: Representative immunohistochemistry with an anti-FLAG antibody on HEK293 cells stably expressing TAP-tagged Na_v_1.7 (green). Right panel: The cell nuclei were co-stained with DAPI (blue). Scale bar = 25 μm. C Representative current traces recorded from HEK293 cells stably expressing either the wild-type Na_v_1.7 or the TAP-tagged Na_v_1.7 in response to depolarisation steps from - 100 to 40 mV. D Left panel: I(V) curves obtained in Na_V_1.7 and TAP-tagged Na_v_1.7 expressing cells using the same protocol as in (c), showing no significant difference in voltage of half-maximal activation (V_1/2_; −9.54 ± 1.08 mV for Na_v_1.7, *n* = 10; and −8.12 ± 1.07 mV for TAP-tagged Na_v_1.7, *n* =14; *p* = 0.3765, Student’s *t*-test) and reversal potential (V_rev_; 61.73 ± 3.35 mV for Na_v_1.7, *n* = 10; and −8.12 ± 1.07 mV for TAP-tagged Na_v_1.7, *n* = 14; *p* = 0.9114, Student’s *t*-test). Right panel: Voltage dependence of fast inactivation, assessed by submitting the cells to a 500 ms prepulse from −130 to −10 mV prior to depolarization at −10 mV. No significant difference in the voltage of half inactivation was observed between the two cell lines (V_1/2_ of-64.58 ± 1.32 mV for Na_v_1.7, *n* =10; and - 64.39 ± 1.06 for TAP-tagged Na_v_1.7, *n* =14; *p* = 0.9100, Student’s *t*-test).

### Generation of a TAP-tagged Na_V_1.7 mouse

We used a conventional gene targeting approach to generate an epitope-tagged Na_V_1.7 mouse. The gene targeting vector was constructed using an *Escherichia coli* recombineering based method (Bence et al., 2005) (Supplemental Fig S1). A TAP tag, which contains a HAT domain, a TEV cleavage site and 3x Flag tags (Fig 1A), was inserted into the open reading frame at the 3’-end prior to the stop codon in exon 27 of Na_V_1.7 (NCBI Reference: NM_001290675) (Fig 2A). The final targeting vector construct containing a 5’ homology arm (3.4 kb), a TAP tag, neomycin cassette and a 3’ homology arm (5.8 kb) (Fig 2B) was transfected into the 129/Sv embryonic stem (ES) cells. 12 colonies with the expected integration (targeting efficiency was 3.5%) were detected using Southern blot (Fig 2C). Germline transmission and intact TAP tag insertion after removal of the neo cassette was confirmed with Southern blot, PCR and RT-PCR, respectively (Fig 2D-F). This mouse line is henceforth referred to as Na_V_1.7^TAP^.

**Figure 2.**
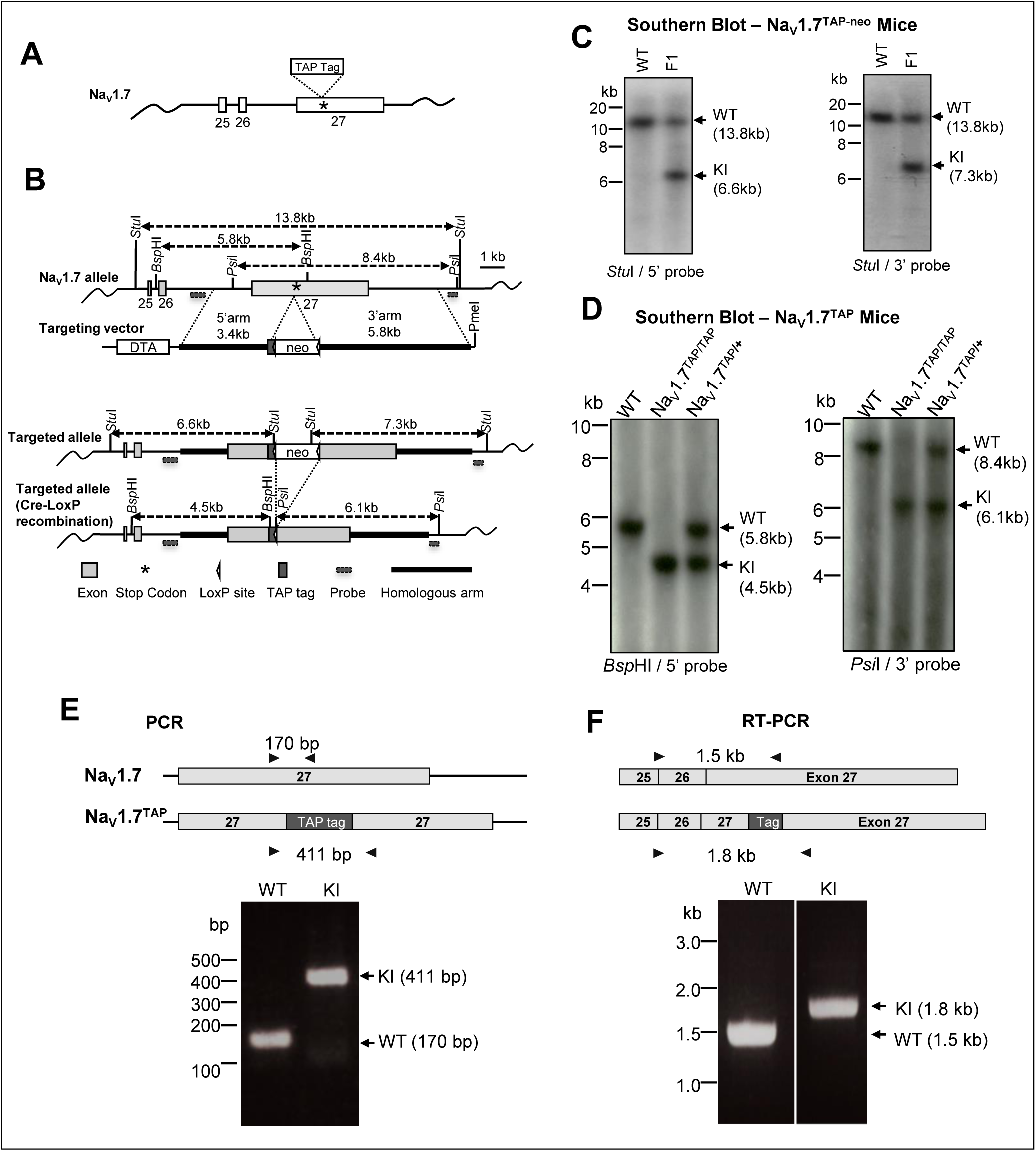
Generation of TAP-tagged Na_V_1.7 knock-in mice. A The location of TAP tag in the Na_V_1.7 locus. A sequence encoding a TAP tag peptide comprised of a HAT domain, TEV cleavage site and 3 FLAG-tags was inserted immediately prior to the stop codon (indicated as a star) at the extreme C-terminus of Na_V_1.7. B Schematic diagrams of the targeting strategy. White boxes represent _V_1.7 exons (exon numbers are indicated on the box), grey box represents TAP tag, thick black lines represent homologous arms, and the small triangle box represents the single LoxP site, respectively. The positions of the external probes used for Southern blotting are indicated in the diagram. Neomycin (neo), DTA expression cassettes and restriction enzyme sites and expected fragment sizes for Southern blotting are also indicated. C Southern blot analysis of genomic DNA from Founder 1 mice (TAP-tagged Na_V_1.7 carrying the neo cassette). Genomic DNA was digested with *Stu*I and was then hybridized with either 5’ or 3’ external probe. Wild-type (WT) allele was detected as a 13.8 kb fragment using either 5’ or 3’ probes. Knock-in allele (KI) was detected as either a 6.6 kb (5’ probe) or a 7.3 kb (3’ probe) fragment. D Southern blot analysis of TAP-tagged Na_V_1.7 allele after Cre recombination. Genomic DNA was digested either with *Bsp*HI or *Psi*l, and was then hybridized with either 5’ (*Bsp*HI) or 3’ (*Psi*l) external probe. WT alleles were detected as 5.8 kb (5’ probe) and 8.4 kb (3’ probe) fragments, respectively. The neo-deleted TAP-tagged Na_V_1.7 alleles were detected as 4.5 kb (5’ probe) and 6.1 kb (3’ probe) fragments, respectively. E Genotyping analysis by PCR. Representative result of the PCR screening of Na_V_1.7^TAP^ mice showing the 411 bp band (KI allele) and the 170 bp band (WT allele). The location of primers used for PCR are indicated with black arrows. F TAP-tagged Na_V_1.7 expression analysis with RT-PCR. Total RNA was isolated from DRG of Na_V_1.7^TAP^ mice and cDNA synthesis was primed using oligo-dT. PCR was performed with the primers indicated with black arrows. A 1.5 kb WT band and a 1.8 kb KI band were detected from either littermate WT control animals or Na_V_1.7^TAP^ KI mice, respectively.

### Na_V_1.7^TAP^ mice have normal pain behavior

The homozygous Na_V_1.7^TAP^ knock-in mice (KI) were healthy, fertile and apparently normal. Motor function of the mice was examined with the Rotarod test. The average time that KI animals stayed on the rod was similar to the WT mice (Fig 3A), suggesting there is no deficit on motor function in the KI mice. The animal responses to low-threshold mechanical, acute noxious thermal, noxious mechanical stimuli, and acute peripheral inflammation were examined with von Frey filaments, Hargreaves’ test, Randall-Selitto apparatus and Formalin test, respectively. The results showed that TAP-tagged KI mice had identical responses to these stimuli compare to the littermate WT control mice (Fig 3B-E), indicating Na_V_1.7^TAP^ mice have normal acute pain behavior.

**Figure 3.**
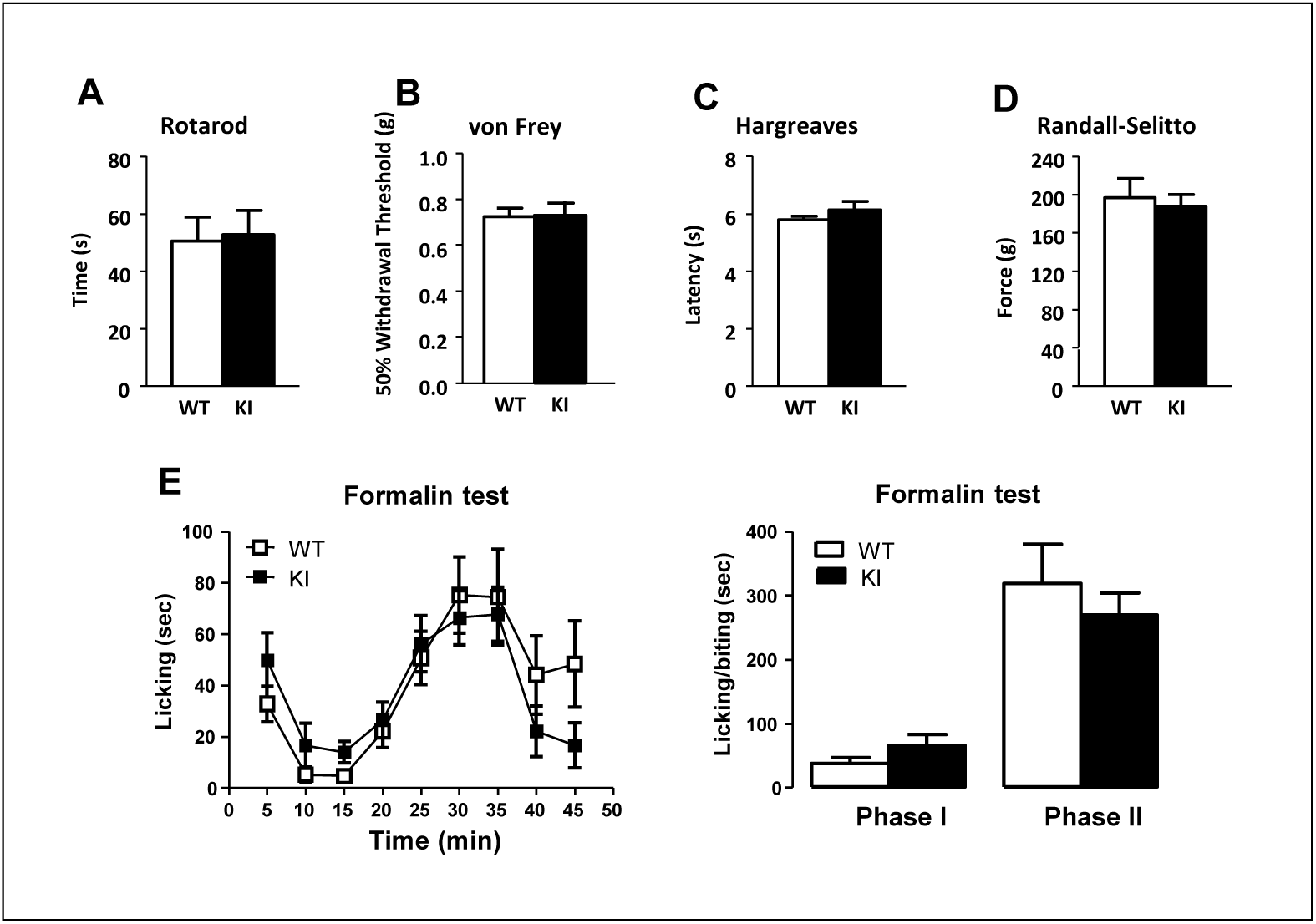
Pain behavior tests. A Rotarod test showed no motor deficits in TAP-tagged Na_V_1.7 animals (n = 7, WT; n = 7, KI; *p* = 0.8619, Student’s *t*-test). B Responses to low-threshold mechanical stimulation by von Frey filaments were normal in the KI mice (*p* = 0.9237, Student’s *t*-test). C Hargreaves’ apparatus demonstrated identical latencies of response to thermal stimulation (n = 7, WT; n = 7, KI; *p* = 0.3038, Student’s *t*-test). D Acute mechanical pressure applied with the Randall-Selitto apparatus demonstrated identical behavior in KI and WT mice (n = 7, WT; n = 7, KI; *p* = 0.7124, Student’s *t*-test). E Formalin test. Licking/biting response to acute peripheral inflammation induced by intraplantar injection of 5% formalin in hind-paw was recorded. Left panel, the time course of development of the response of KI mice (black squares) and WT littermate controls (white squares) showed similar response patterns (n = 10, WT; n = 7, KI; *p* = 0.2226, two-way ANOVA). Right panel, the early (0 - 10 minutes) and late (10 - 45 minutes) phases of the formalin response in KI and WT mice showed similar responses (*p* = 0.1690, 1^st^ phase; *p* = 0.5017, 2^nd^ phase; Student’s *t*-test) between KI and WT mice.

### Expression pattern of TAP-tagged Na_V_1.7 in the nervous system

We used immunohistochemistry to examine the expression pattern of TAP-tagged Na_V_1.7 in the nervous system of Na_V_1.7^TAP^ mice. The results showed that the FLAG-tag was expressed in the olfactory bulb, with strong staining visible in the olfactory nerve layer, the glomerular layer, and in the accessory olfactory bulbs (Fig 4A-D), consistent with previous results (Weiss et al., 2011). In the brain, FLAG-tag expression was present in the medial habenula, the anterodorsal thalamic nucleus, the laterodorsal thalamic nucleus, and in the subfornical organ that is located in the roof of the dorsal third ventricle (Fig 4E,F) and is involved in the control of thirst (Oka et al., 2015). The FLAG-tag was also present in neurons of the posterodorsal aspect of the medial amygdala and in the hypothalamus in neurons of the arcuate nucleus (Fig 4G-I) as confirmed in a recent study (Branco et al., 2016). A clear staining appeared in neurons of the substantia nigra reticular region and the red nucleus magnocellular region of the midbrain, and in neurons of the pontine nuclei located in the hindbrain (Fig 4J-L). In the spinal cord, FLAG-tag expression was visible in the superficial layer of the dorsal horn. We co-stained the spinal cord with IB4, a marker for the inner part of lamina II, and the results showed that the TAP-tagged Na_V_1.7 was expressed in Laminae I, II and III (Fig 4M-O). In the PNS, however, there was no positive FLAG-tag staining found in DRG, sciatic nerve or skin nerve terminals (data not shown). This could be because of masking of the tag in the PNS, preventing the antibody binding. We also examined the TAP-tagged Na_V_1.7 expression pattern in different tissues with Western blot using an anti-FLAG antibody. TAP-tagged Na_V_1.7 bands were present in olfactory bulb, hypothalamus, spinal cord, sciatic nerve and DRG, but not in cortex, cerebellum, skin, lung, heart and pancreas (Fig 5E).

**Figure 4.**
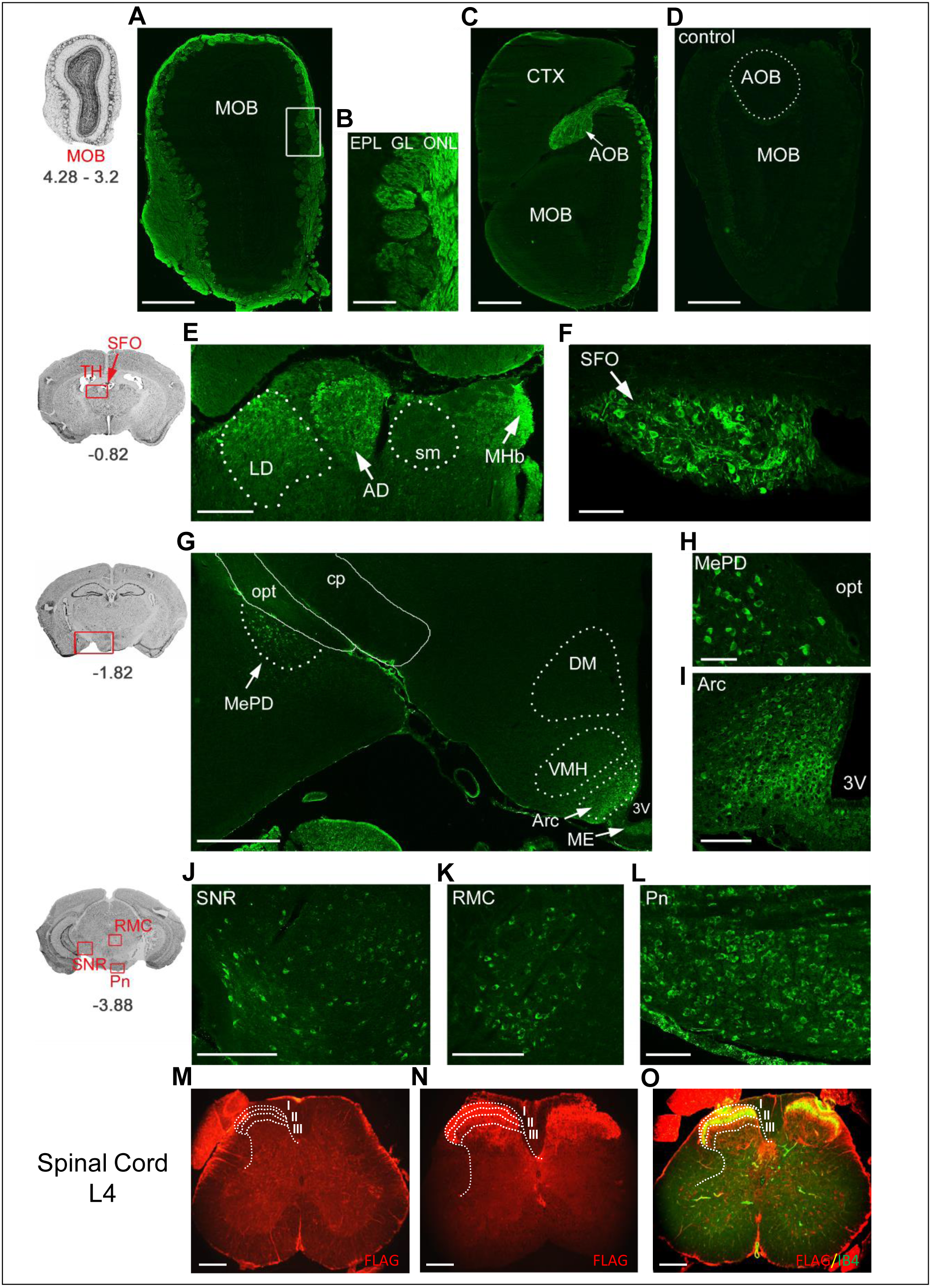
Immunohistochemistry of FLAG-tag expression in the central nervous system (CNS). A-D In the main olfactory bulb (MOB), FLAG-tag expression (in green) is visible in the olfactory nerve layer (ONL) and in the glomerular layer (GL) in TAP-tagged Na_V_1.7 knock-in mice (KI) but not in the littermate wild-type controls (WT). In the posterior olfactory bulb, staining is also evident in the (C) accessory olfactory bulb (AOB). Staining is absent in the (D) MOB and AOB of wild-type control mice. E-F FLAG-tag expression is present in the medial habenula (MHb, arrow), the anterodorsal thalamic nucleus (AD, arrow), the laterodorsal thalamic nucleus (LD, dotted line), and (F) in the subfornical organ (SFO, arrow) located in the roof of the dorsal third ventricle. G-I FLAG-tag expression is present in neurons of the posterodorsal aspect of the medial amygdala (MePD, arrow, dotted line) and in the hypothalamus in neurons of the arcuate nucleus (Arc, arrow, dotted line). J-L FLAG-tag expression is present (J) in neurons of the substantia nigra reticular part (SNR) and (K) the red nucleus magnocellular part (RMC) of the midbrain, and (L) in neurons of the pontine nuclei (Pn) located in the hindbrain. cp, cerebral peduncle; CTX, cortex; DM, dorsomedial hypothalamic nucleus; EPL, external plexiform layer; ME, median eminence; opt, optic tract; sm, stria medullaris; TH, thalamus; VMH, ventromedial hypothalamic nucleus; 3V, third ventricle. M-N The cross section of lumbar spinal cord (L4) is labelled with anti-FLAG (in red). FLAG tag expression is present in laminae I, II and III in spinal cord of KI mice (N) but not in spinal cord of WT mice (M). O The cross sections of spinal cord of KI mice were co-stained with Laminae II marker IB4 (o, in green). Sketches on the left illustrate the CNS regions and bregma levels (in mm) of the fluorescence images shown on the right. Scale bars: 500 μm (A, C, D, G); 250 μm (E, J, K, M-O); 100 μm (B, F, I, L); 50 μm (H).

### Optimisation of single- and tandem-affinity purification of TAP-tagged Na_V_1.7

The TAP tag on Na_V_1.7 offered a possibility of two consecutive affinity purifications, a single-step affinity purification (ss-AP) (Step 1 and 2) and tandem affinity purification (Step 1 - 4) (Fig 5A). CHAPS and DOC lysis buffers were evaluated to solubilize TAP-tagged Na_V_1.7 and its protein complex from tissues, e.g. DRG, spinal cord, olfactory bulbs and hypothalamus, in the Co-IP system. CHAPS is a non-denaturating zwitterionic detergent, commonly used to extract membrane proteins in their native conformation. In comparison to strong anionic detergents like SDS, CHAPS preserves protein-protein interactions and is compatible with downstream applications such as mass spectrometry. The result showed that TAP-tagged Na_V_1.7 from DRG and olfactory bulbs was clearly solubilized and precipitated by both purifications – ss-AP and tandem affinity purification in 1% CHAPS buffer (Fig 5B). Also the result from ss-AP showed that TAP-tagged Na_V_1.7 could be immunoprecipitated from hypothalamus, sciatic nerve, spinal cord, olfactory bulb and DRG of KI mice, but not from these tissues of WT control mice (Fig 5C). However, the DOC lysis buffer, which was used to investigate the TAP-tagged PSD-95 protein complex (Fernandez et al., 2009), did not solubilize TAP-tagged Na_V_1.7 from mouse tissue (data not shown).

**Figure 5.**
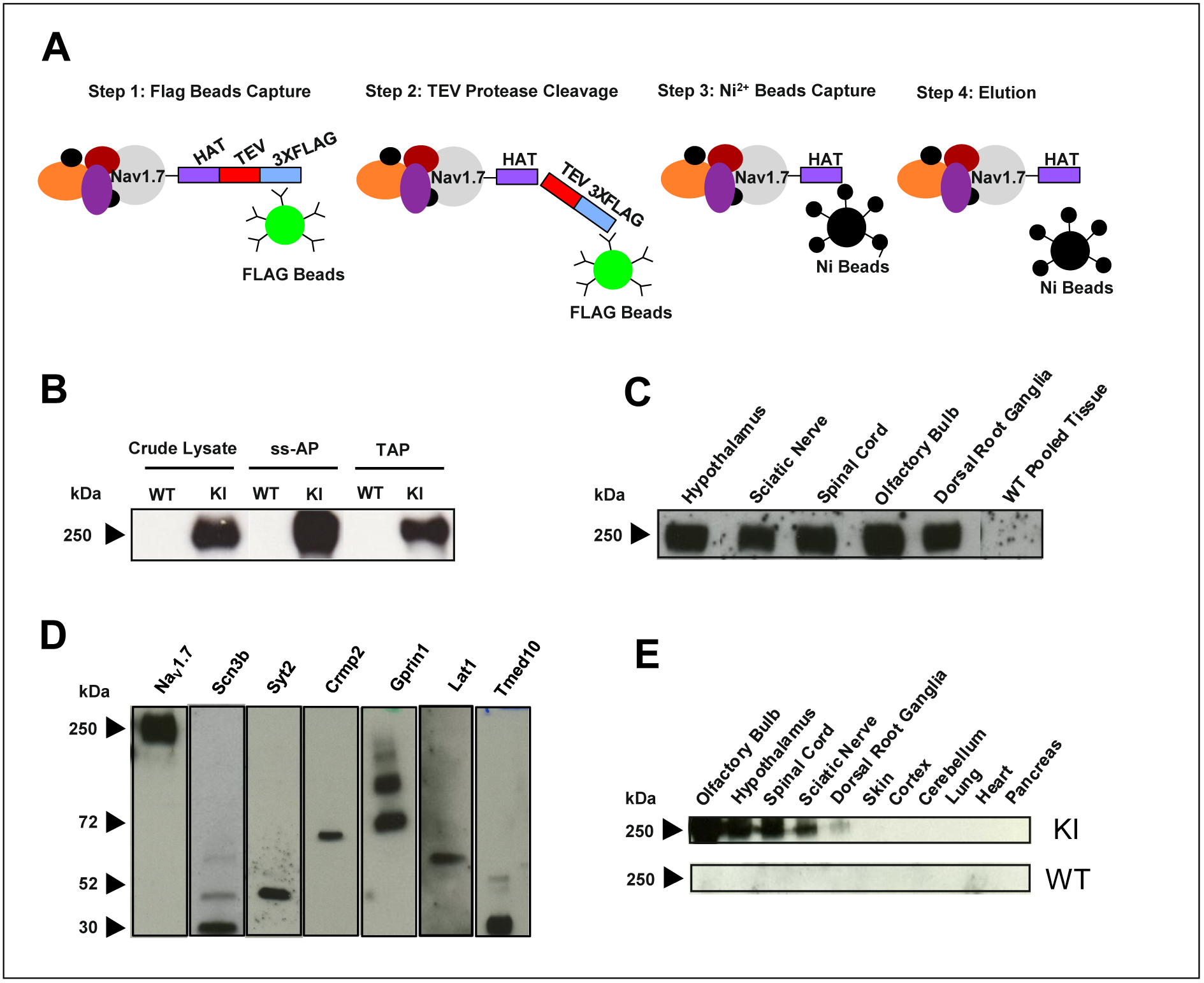
Optimisation of single-step and tandem affinity purification, validation of identified protein-protein interactors and tissue expression pattern of TAP-tagged Na_V_1.7 in Na_V_1.7^TAP^ mice. A Schematic illustrating the affinity purification (ss-AP and TAP) procedure using the tandem affinity tags separated with a TEV cleavage site. B The proteins from DRG and olfactory bulbs were extracted in 1*%* CHAPS lysis buffer. After single-step and tandem affinity purification, TAP-tagged Na_V_1.7 were detected using Western blotting with anti-HAT antibody. C The proteins from different tissues including hypothalamus, sciatic nerve, spinal cord, olfactory bulbs and DRG from KI mice, and pooled tissues from WT mice were extracted in 1% CHAPS lysis buffer. After single-step affinity purification, TAP-tagged Na_V_1.7 was detected using Western blotting with anti-HAT antibody. D The interaction between TAP-tagged Na_V_1.7 and identified Na_V_1.7 protein-protein interactors including Scn3b, Syt2, Crmp2, Gprin1, Lat1 and Tmed10 were validated using a co-immunoprecipitation *in vitro* system. The expression vectors containing cDNA of validated genes were cloned and transfected into a HEK293 cell line stably expressing TAP-tagged Na_V_1.7. After transfection, TAP-tagged Na_V_1.7 complexes were immunoprecipitated with anti-FLAG antibody and the selected candidates were detected with their specific antibody using Western blotting. The results showed the expected sizes of Scn3b (32 kDa), Syt2 (44 kDa), Crmp2 (70 kDa), Gprin1 (two isoforms: 80 kDa and 110kDa), Lat1 (57 kDa) and Tmed10 (21 kDa). E Tissue expression pattern of TAP-tagged Na_V_1.7. The proteins were extracted from different tissues in both KI and WT littermate control mice and anti-FLAG used to detect TAP-tagged Na_V_1.7 using western blotting.

### Identification of TAP-tagged Na_V_1.7 associated complexes by AP-MS

We next identified the components of Na_V_1.7 complexes using ss-AP followed by LC-MS/MS. Briefly, the TAP-tagged Na_V_1.7 complexes were extracted from DRG, spinal cord, olfactory bulb and hypothalamus using ss-AP (see Materials and Methods). In total 189,606 acquired spectra from 12 samples, in which each group (KI and WT) contains 6 biological replicate samples and each sample was from one mouse, were used for protein identification. 1,252 proteins were identified with a calculated 0.96% false discovery rate (FDR). 267 proteins (Supplemental Tab S1) met those criteria and were shortlisted based on the criteria described in Materials and Methods. The proteins only appearing in Na_V_1.7^TAP^ mice and the representatively selected proteins are listed in Table 1. PANTHER cellular component analysis of these 267 proteins revealed 8 different cellular components (Fig 6A). These proteins were further classified into 22 groups based on their function, including 12 membrane trafficking proteins, 23 enzyme modulators and 4 transcription factors (Fig 6B).

**Table 1.**
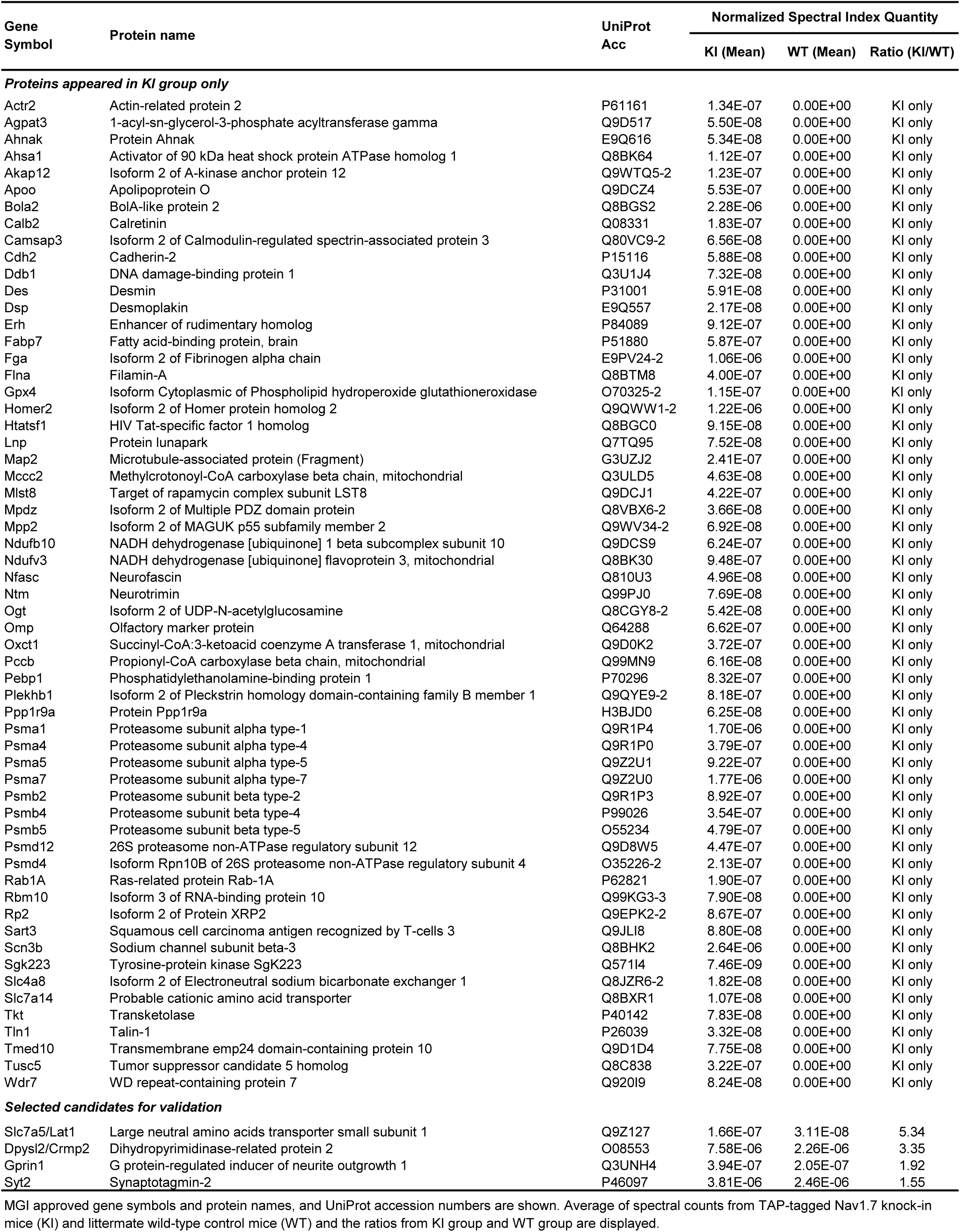
Identified Na_V_1.7 associated proteins (only appeared in KI group + selected candidates)

**Figure 6.**
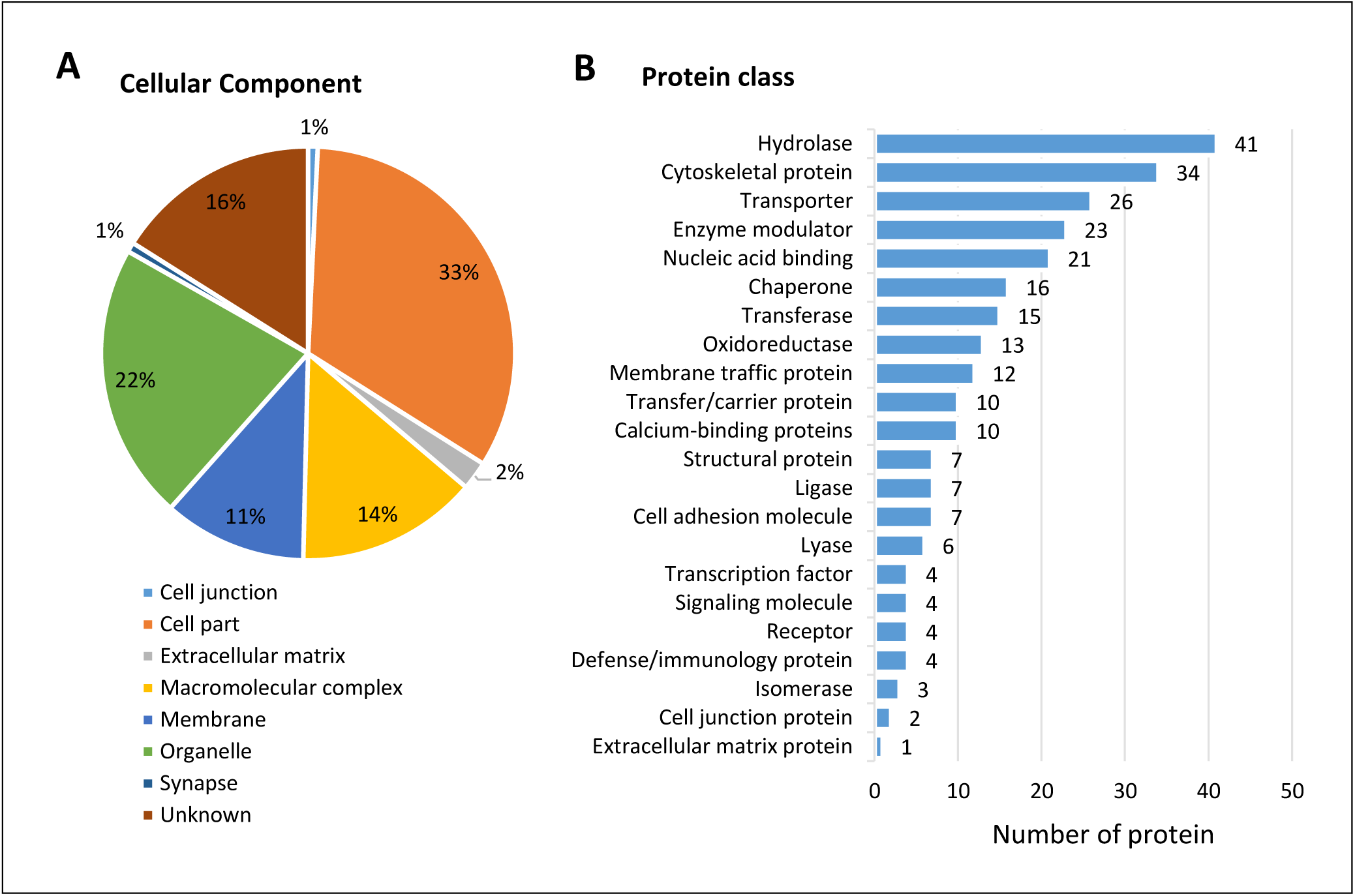
Analysis of Na_V_1.7 complex proteins. A Cellular localization of identified Na_V_1.7 interacting proteins. B Protein class of identified Na_V_1.7 interacting proteins categorized using PANTHER Classification System.

### Validation of TAP-tagged Na_V_1.7 interacting proteins using Co-IP

The physical interactions between Na_V_1.7 and interacting protein candidates were assessed using Co-IP in an *in vitro* system by co-expressing the candidates in a TAP-tagged Na_V_1.7 stable cell line. A number of candidates of interest, such as Syt2, Gprin1, Lat1, and Tmed10, together with known Na_V_1.7 protein interactors Scn3b and Crmp2 were chosen from the identified Na_V_1.7 associated protein list (Supplemental Tab S1 and Table 1) for further validation.

Scn3b and Crmp2 have previously been implied to associate with Na_V_1.7 (Dustrude et al., 2013, Ho et al., 2012) and firstly we confirmed the physical interaction between these proteins and Na_V_1.7 by co-immunoprecipitation (Fig 5D). Next, we investigated some potential novel binding partners, such as Syt2, Gprin1, Lat1, and Tmed10. Synaptotagmin is a synaptic vesicle membrane protein that functions as a calcium sensor for neurotransmission (Chapman, 2002). Sampo *et al.* showed a direct physical binding of synaptotagmin-1 with Na_V_1.2 at a site which is highly conserved across all voltage-gated sodium channels (Sampo et al., 2000) suggesting that the synaptotagmin family can associate with other VGSCs. Na_V_1.7 found at pre-synaptic terminals appears to be involved in neurotransmitter release (Minett et al., 2012, Black et al., 2012). Our Co-IP result shows that Syt2 was co-precipitated with TAP-tagged Na_V_1.7 (Fig 5D). Interestingly other synaptic proteins also co-precipitated with Na_v_1.7 including SNARE complex protein Syntaxin-12 (Supplemental Tab S1). Gprin1 was selected for further validation due to its involvement in regulating mu-opioid receptor (MOR) activity (Ge et al., 2009), highlighting a potential role in connection between the opioid system and Na_V_1.7 in pain. Our Co-IP results confirmed the protein interaction between Gprin1 and Na_V_1.7 (Fig 5D). Gabapentin was developed to treat epilepsy, but it is now used to treat various forms of chronic pain. However, the analgesic mechanisms of gabapentin are not entirely clear, but involve inhibiting the function of N-type voltage-gated calcium channels (CaV2.2) by blocking the association of pore-forming α1 subunits with auxiliary α2δ-1 subunits to reduce channel activity, membrane trafficking, and neurotransmitter release in DRG neurons (Kukkar et al., 2013). As the transporter of gabapentin, Lat1 has been confirmed with Co-IP (Fig 5D), suggesting that Lat1 and/or gabapentin may be involved in the regulation of Na_V_1.7 channel function. However, future experiments are required to determine the functional significance of the Na_v_1.7-Lat1 interaction in this regard. As a member of the p24 family, Tmed10 was selected due to its well-characterised properties as a protein trafficking regulator. Previous studies have highlighted that Tmed10 plays an important role as a cargo receptor in the trafficking of various membrane proteins. For example, Tmed10 has been observed to modulate the transport and trafficking of amyloid-β precursor protein (Vetrivel et al., 2007), endogenous glycosylphosphatidylinositol (GPI)-anchored proteins CD59 and folate receptor alpha (Bonnon et al., 2010), and several G protein-coupled receptors (GPCRs) (Luo et al., 2011). Our Co-IP result showed that Tmed10 was co-immunoprecipitated with TAP-tagged Na_V_1.7 (Fig 5D), suggesting Tmed10 may regulate Na_V_1.7 trafficking. In summary, all these four potential interacting proteins selected for further validation were confirmed as interactors by Co-IP, giving confidence in the Na_v_1.7^TAP^ mass spectrometry list.

### Functional characterisation of Crmp2

Crmp2 is the presumed target of the anti-epileptic drug Lacosamide based on binding studies (Wilson and Khanna, 2015). We sought to evaluate the possible electrophysiological effects of Crmp2 on Na_V_1.7 and in addition understand the relevance of Crmp2 to the action of the drug on Na_V_1.7 current density. Transfection of Crmp2 into a Na_V_1.7 stable HEK293 cell line revealed a nearly two-fold increase in sodium current density (Fig 7A-C), suggesting Crmp2 acts as a transporter for Na_V_1.7 (Dustrude et al., 2013). Next, we sought to investigate the effect of Lacosamide on sodium currents. In cells not transfected with Crmp2, Na_V_1.7 currents displayed no significant changes in current density following prolonged 5-hour exposure to Lacosamide (Fig 7A,B). Interestingly, following 5-hour incubation with Lacosamide in Crmp2 transfected cells, a complete reversal in the increase in Na_V_1.7 current density provoked by Crmp2 was observed (Fig 7A,D).

**Figure 7.**
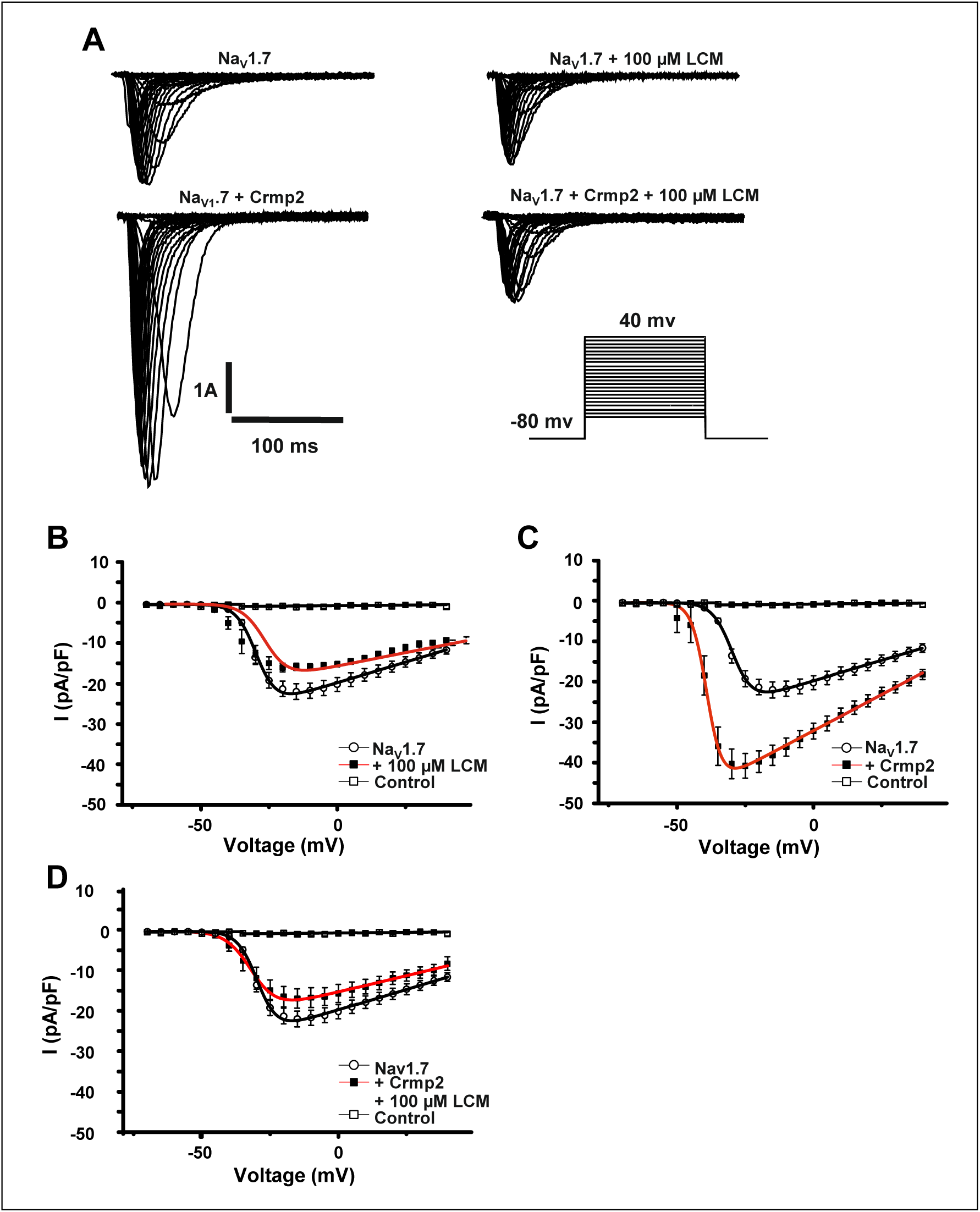
Electrophysiological characterisation of Na_V_1.7 in HEK293 cells following transfection with Crmp2 and incubation with Lacosamide (LCM). A Representative raw current traces of Na_V_1.7 stably expressed in HEK293 cells in response to the activation pulse protocol shown. Each trace shows a different condition: + Crmp2 transfection and + incubation with 100 μΜ LCM. B IV plot of Na_V_1.7 current denity in the absence and presence of 100 μΜ LCM. Compared with Na_V_1.7 basal currents (n = 10), LCM incubation had no significant effect on sodium channel density (n = 12, *p = 0.402*). C IV plot of Na_V_1.7 in HEK293 cells in the presence and absence of Crmp2 transfection. Compared to Na_V_1.7 basal currents, Crmp2 transfection caused a significant increase in Na_V_1.7 current density (n = 16, *p* = 0.0041). D IV plot showing current density of Na_V_1.7 following transfection of Crmp2 and incubation with 100 μΜ LCM. Incubation with LCM reversed the Crmp2-mediated current increase (n = 10, *p* = 0.0442). Data were analysed using one-way ANOvA with Tukey’s *post hoc* test.

## Discussion

Experimental evidence has shown that protein-protein interactions play a key role in trafficking, distribution, modulation and stability of ion channels (Catterall, 2010, Bao, 2015, Chen-Izu et al., 2015, Laedermann et al., 2015, Shao et al., 2009, Leterrier et al., 2010). Here, we mapped the protein-interaction network of Na_V_1.7 using an AP-MS proteomic approach with an epitope-tagged Na_V_1.7 knock-in mouse line. This is the first report to define an ion channels’ macromolecular complex using an epitope-tagged gene targeted mouse.

AP-MS requires specific high-affinity antibodies against the target proteins of interest (Wildburger et al., 2015). However, binding may compete with normal protein-protein interactions. To overcome these limitations, epitope tags on target proteins were introduced into the AP-MS system. In the last decade, single-step and tandem affinity purification have been widely applied in protein-protein interaction studies (Wildburger et al., 2015, Fernandez et al., 2009). In contrast to ss-AP, TAP produces lower background and less contamination. However, due to its longer experimental washing procedures and two-step purification, TAP coupled with MS analysis may not be a sensitive method to detect transient and dynamic protein-protein interactions. In recent years, along with newly developed highly sensitive mass spectrometer techniques and powerful quantitative proteomics analysis methods, ss-AP was employed to identify both transient and stable protein-protein interactors (Keilhauer et al., 2015, Oeffinger, 2012). For example, using a single-step FLAG approach, Chen and colleagues defined specific novel interactors for the catalytic subunit of PP4, which they had not previously observed with TAP-MS (Chen and Gingras, 2007). Thus, ss-AP followed by sensitive LC-MS/MS analysis was applied in this study. In fact, many dynamic modulator proteins were identified to interact with Na_V_1.7 in this study, such as Calmodulin (Calm1) (Supplemental Tab S1) which was found to bind to the C-terminus of other VGSCs Na_V_1.4 and Na_V_1.6, and regulates channel function (Herzog et al.,2003). Four proteins, Crmp2, Nedd4-2, FGF13 and Pdzd2, have previously been reported as Na_V_1.7 protein interactors (Ho et al., 2012, Sheets et al., 2006, Shao et al., 2009, Yang et al., 2017). Laedermann *et al.* showed that the E3 ubiquitin ligase Nedd4-2 regulates Na_V_1.7 by ubiquitination in neuropathic pain (Laedermann et al., 2013). Shao *et al.* demonstrated that Pdzd2 binds to the second intracellular loop of Na_V_1.7 by a GST pull-down assay in an *in vitro* system (Shao et al., 2009). We did not find the previously reported Na_V_1.7 interactors Nedd4-2, FGF13 and Pdzd2. This may be because Nedd4-2 and FGF13 only binds to Na_V_1.7 in neuropathic pain and heat nociception conditions, respectively, and Pdzd2 shows strong binding *in vitro* but not *in vivo.* Apart from known Na_V_1.7 and other Na_V_ protein interactors, we also identified a broad range of important novel interactors that belong to different protein classes, such as cytoskeletal/structural/cell adhesion proteins and vesicular/trafficking/transport proteins (Fig 6B).

Using Co-IP and a co-expression *in vitro* system, we confirmed a direct physical interaction between Na_V_1.7 and novel Na_V_1.7 interactors, such as Syt2, Gprin1, Lat1 and Tmed10, and known interactors Scn3b and Crmp2 as well. Furthermore, we demonstrated that transient over-expression of Crmp2 can up-regulate Na_V_1.7 current density in stably expressing Na_V_1.7 HEK293 cells and this up-regulation can be reversed by applying Lacosamide. Taken together, FLAG ss-AP coupled with quantitative MS seems to be a powerful and reliable tool for investigating protein interactions of membrane ion channels.

VGSCs are known to exist in macromolecular complexes (Meadows and Isom, 2005). The β subunits are members of the immunoglobulin (Ig) domain family of cell-adhesion molecules (CAM). As well as sodium channel α subunits, the β subunits also bind to a variety of cell adhesion molecules such as Neurofascin, Contactin, Tenascins, and NrCAMs (Srinivasan et al., 1998, McEwen and Isom, 2004, Ratcliffe et al., 2001, Cusdin et al., 2008, Namadurai et al., 2015). In our data set (Supplemental Tab S1), the sodium channel β3 subunit and some CAMs such as Ncam1 and Neurofascin, have been found to associate with the Na_V_1.7 α subunit. The crystal structure of the human β3 subunit has been solved recently. The β3 subunit Ig-domain assembles as a trimer in the crystal asymmetric unit (Namadurai et al., 2014). This raises the possibility that trimeric β3 subunits binding to Na_V_1.7 α subunit(s) form a large complex together with other sodium channels, as well as CAMs and cytoskeletal proteins in the plasma membrane.

Na_V_1.7 has also been linked to opioid peptide expression, and enhanced activity of opioid receptors is found in the Na_V_1.7 null mutant mouse (Minett et al., 2015). Interestingly, Gprin1, which is known to interact with opioid receptors as well as other GPCRs, was found to co-immunoprecipitate with Na_V_1.7. This suggests that GPCR sodium channel interactions could add another level of regulatory activity to the expression of Na_V_1.7. More recently, Branco and colleagues reported that Na_V_1.7 in hypothalamic neurons plays an important role in body weight control (Branco et al., (2016). We found that Na_V_1.7 was not only present in the arcuate nucleus but also in other regions of the brain such as the medial amygdala, medial habenula, anterodorsal thalamic nucleus, laterodorsal thalamic nucleus, and in the subfornical organ, substantia nigra reticular part and the red nucleus magnocellular part of the midbrain, and in neurons of the pontine nuclei located in the hindbrain. Na_V_1.7 thus has other functions in the CNS that remain to be elucidated.

Overall, the present findings provide new insights into the interactome of Na_V_1.7 for advancing our understanding of Na_V_1.7 function. Our data also show that the ss-AP coupled LC-MS/MS is a sensitive, reliable and high-throughput approach to identify protein-protein interactors for membrane ion channels, using epitope-tagged gene targeted mice.

## Materials and Methods

### Generation of a TAP-tagged Na_V_1.7 expressing stable HEK293 cell line

A HEK293 cell line stably expressing TAP tagged Na_V_1.7 was established as previously described (Koenig et al., 2015). Briefly, a sequence encoding a TAP tag (peptide: SRK DHL IHN VHK EEH AHA HNK IEN LYF QGE LPT AAD YKD HDG DYK DHD IDY KDD DDK) was inserted immediately prior to the stop codon of Na_V_1.7 in the SCN9A mammalian expression construct FLB (Cox et al., 2006). The TAP tag at the extreme C-terminus of Na_V_1.7 comprises a HAT domain and 3 FLAG tags, enabling immunodetection with either anti-HAT or anti-FLAG antibodies. The function and expression of TAP tagged Na_V_1.7 in this HEK293 cell line were characterized with both immunocytochemistry and electrophysiological patch clamp analysis.

### Generation of Na_V_1.7^TAP^ knock-in mice

The gene targeting vector was generated using a BAC homologous recombineering-based method (Liu et al., 2003). Four steps were involved in this procedure (Fig 2 – figure supplement 1). Step 1, two short homology arms (HA) HA3 and HA4 corresponding to 509 bp and 589 bp sequences within intron 26 and after exon 27 of Na_V_1.7, respectively, were amplified by PCR using a BAC bMQ277g11 (Source Bioscience, Cambridge UK) DNA as a template, and then inserted into a retrieval vector pTargeter (Fernandez et al., 2009) (a gift from Dr Seth GN Grant) by subcloning. Step 2, a 9.1 kb genomic DNA fragment (3.4 kb plus 5.8 kb) was retrieved through homologous recombineering by transforming the *KpnI*-linearized pTargeter-HA3-HA4 vector into EL250 *E. coli* cells containing BAC bMQ277g11. Step 3, short homology arms HA1 and HA2 corresponding to 550 bp (before stop codon of Na_V_1.7) and 509 bp (starting from stop codon of Na_V_1.7) respectively were amplified by PCR, and then cloned into pneoflox vector (Fernandez et al., 2009) (a gift from Dr Seth GN Grant) containing TAP tag, leaving in between the TAP tag sequence, 2 LoxP sites, PGK and EM7 promoters, the G418^r^ gene and a SV40 polyadenylation site. Step 4, the cassette flanked by two homology arms (HA1 and HA2) was excised by *XhoI* and *Bgl*II digestion and transformed into recombination-competent EL250 cells containing pTargeter-HA3-HA4 plasmid. Then, the TAG tag cassette was inserted into the pTargeter-HA3-HA4 vector by recombination in EL250 *E. coli* cells. The correct recombination and insertion of the targeting cassette were confirmed by restriction mapping and DNA sequencing. The complete gene targeting vector containing 5’-end homologous Na_V_1.7 sequence of 3.4 kb and a 3’-end homology arm of 5.8 kb was linearized with *PmeI* digestion for ES cell electroporation. All the homology arms HA1, HA2, HA3 and HA4 were amplified with NEB Phusion PCR Kit using bMQ277g111 BAC clone DNA as a template. Primers used to create the recombination arms included:

HA1XhoIF (HA1, forward) - acactcgagAGCCAAACAAAGTCCAGCT
HA1XbaIR (HA1, reverse) - tgttctagaTTTCCTGCTTTCGTCTTTCTC
HA2Acc65IF (HA2, forward) - tgaggtacctagAGCTTCGGTTTTGATACACT
HA2BgIIIR (HA2, reverse) - gatagatctTTGATTTTGATGCAATGTAGGA
HA3SpeIF (HA3, forward) - ctcactagtCTCTTCATACCCAACATGCCTA
HA3KpnIR (HA3, reverse) - aatggtaccGGATGGTCTGGGACTCCATA
HA4KpnIF (HA4, forward) - gaaggtaccGCTAAGGGGTCCCAAATTGT
HA4PmeIR (HA4, reverse) - tcagtttaaacGGGATGGGAGATTACGAGGT

To generate TAP tagged Na_V_1.7 mice, the linearized targeting vector was transfected into 129/Sv ES cells. Cells resistant to G418 were selected by culturing for 9 days. Recombined ES cell clones were identified using Southern blot-based screening. Three Clones that were confirmed to be correct using Southern blot were injected into C57BL/6 blastocysts at the Transgenic Mouse Core facility of the Institute of Child Heath (ICH). The chimeric animals were crossed to C57BL/6 and the germline transmission was confirmed by Southern blot. The neomycin cassette was removed by crossing with global Cre mice. The correct removal of the neomycin cassette and TAP tag insertion was confirmed by Southern blot, genotyping (PCR) and RT-PCR.

The genomic DNA was extracted from ear punch and PCR genotyping was performed as described previously (Dicer 2008). Primers used for PCR included:

5aF1 (forward) – ACAGCCTCTACCATCTCTCCACC
3aR4 (reverse) - AACACGAGTGAGTCACCTTCGC

The wild-type Na_V_1.7 allele and TAP tagged Na_V_1.7 allele gave a 170bp band and a 411 bp band, respectively. The TAP tagged Na_V_1.7 mRNA was confirmed by RT-PCR. Briefly, the total RNA was extracted from dorsal root ganglia with Qiagen RNAEasy kit (Qiagen) and 1.0 μg of total RNA was used to synthesize cDNA using Biorad cDNA script-II synthesize kit with oligo-dT primer. The following primers were used to detect mRNA of Na_V_1.7:

E25-26-F (forward) – CCGAGGCCAGGGAACAAATTCC
3’UTR-R (reverse) - GCCTGCGAAGGTGACTCACTCGTG

The wild-type and TAP tagged Na_V_1.7 alleles gave a 1521 bp band and a 1723 bp band, respectively.

### Immunocytochemistry

TAP tagged Na_V_1.7-HEK293 cells were plated on poly-D-lysine coated coverslips in 24-well plates and cultured at 37°C/5% CO_2_ in DMEM supplemented with 10% foetal bovine serum (Life Technologies), 50 U/ml penicillin, 50 μg/ml streptomycin and 0.2 mg/ml G418. 24 hours later, the cells were fixed in cooled methanol at −20°C for 10 minutes, and then permeabilised with cooled acetone at −20°C for 1 minute. After 3 washes with 1x PBS, the cells were incubated with blocking buffer containing 1x PBS, 0.3% Triton X-100 and 10% goat serum at room temperature for 30 minutes. Then the fixed cells were incubated with anti-FLAG antibody (1:500 in blocking buffer, F1804, Sigma) at 4°C overnight. After 3 washes with 1x PBS, the cells were incubated with secondary antibody goat anti-mouse IgG conjugated with Alexa Fluor 488 (A11017, Invitrogen) at room temperature for 2 hours. Then the coverslips were washed 3 times in 1x PBS. The cells were mounted with VECTASHIELD HardSet Antifade Mounting Medium with DAPI (H-1400, Vectorlabs) and visualized using a fluorescence Leica microscope.

### Immunohistochemistry

Following anesthesia, mice were transcardially perfused with PBS, followed by 1% (w/v) paraformaldehyde in PBS. Brains and spinal cord were dissected and incubated in fixative for 4 hours at 4°C, followed by 30% sucrose in PBS for 2 days at 4°C. Tissue was embedded in O.C.T. (Tissue-Tek) and snap-frozen in a dry-ice/2-methylbutane bath. Brain coronal and spinal cord cross cryosections (20 μm) were collected on glass slides (Superfrost Plus, Polyscience) and stored at −80°C until further processing. For Flag-tag immunohistochemistry, sections were incubated in blocking solution (4% horse serum, 0.3% Triton X-100 in PBS) for 1 hour at room temperature, followed by incubation in mouse anti-Flag antibody (F-1804, Sigma) diluted 1:200 in blocking solution for 2 days at 4°C. Following three washes in PBS, bound antibody was visualized using either an Alexa 488 or 594 conjugated goat anti mouse secondary antibody (1:800, Invitrogen). The IB4 staining in spinal cord has been described in a previous study (Zhao et al., 2010). To facilitate the identification of brain regions and the corresponding bregma levels, nuclei were counterstained with Hoechst 33342 (1:10,000, Invitrogen). Sections were mounted with a fluorescence mounting medium (DAKO). Fluorescence images were acquired on an epifluorescence microscope (BX61 attached to a DP71 camera, Olympus) or a confocal laser scanning microscope (LSM 780, Zeiss). Images were assembled and minimally adjusted in brightness and contrast using Adobe Photoshop Elements 10. Bregma levels and brain regions were identified according to the stereotaxic coordinates in “The Mouse Brain” atlas by Paxinos and Franklin 2001.

### Behavioral analysis

All behavioral tests were approved by the United Kingdom Home Office Animals (Scientific Procedures) Act 1986. Age (6-12 weeks) -matched KI mice (4 males and 3 females) and littermate wild-type (WT) controls (3 males and 4 females) were used for acute pain behaviour studies. The experimenters were blind to the genetic status of test animals. The Rotarod, Hargreaves’, von Frey and Randall-Selitto tests were performed as described (Zhao et al., 2006). The Formalin test was carried by intraplantar injections of 20 μl of 5% formalin. The mice were observed for 45 minutes and the time spend biting and licking the injected paw were monitored and counted. Two phases were categorised, the first phase lasting 0 - 10 minutes and the second phase 10 - 45 min. All data are presented as mean ± SEM.

### Single-step and tandem affinity purification

In each round of sample preparation for single-step and tandem affinity purification, DRG, olfactory bulbs, spinal cord and hypothalamus samples were homogenized (Precellys ceramic kit 4.1, Peqlab) in 1% CHAPS (30 mM Tris-HCl pH 7.5; 150 mM NaCl, 1% CHAPS, 1 complete EDTA free Protease inhibitor cocktail (Roche) in 10 ml of CHAPS buffer) and further homogenized using an insulin syringe. The lysates were incubated shaking horizontally on ice and clarified by centrifugation at 14,000 g for 8 min at 4°C. Protein concentrations were measured using the BCA Protein Assay Kit (Pierce) and a total starting amount of 10 mg of protein containing supernatant was incubated with magnetic M-270 Epoxy Dynabeads (14311D, Invitrogen) covalently coupled to mouse Anti-Flag M2 (F-1804, Sigma). The coupling was carried out for 2 hours at 4°C using an end-over-end shaker. Magnetic Dynabeads were collected on a DynaMAG rack (Invitrogen) and washed three times in 1% CHAPS Buffer and 1x AcTEV protease cleavage buffer (50 mM Tris-HCl pH 8.0, 0.5 mM EDTA, 1 mM DTT, Invitrogen). Dynabead-captured Na_V_1.7 TAP-tag complex was released from the beads by incubation with AcTEV protease (Invitrogen) for 3 hours at 30^o^C, finalizing the single step purification. For the tandem affinity purification, protein eluates were collected after AcTEV cleavage and 15x diluted in protein binding buffer (50 mM Sodium phosphate, 300 mM NaCl, 10 mM imidazole, 0.01% Tween; pH 8.0). Na_V_1.7 TAP tag was captured using Ni-NTA beads (Qiagen). Ni-NTA beads were washed three times in protein binding buffer and incubated overnight at 4^o^C on an end-over-end shaker. TAP-tagged Na_V_1.7 protein complexes were released from the Ni-NTA beads by boiling in 1x SDS protein sample buffer.

### Western blot

Proteins were extracted from different tissues, such as DRG, spinal cord, olfactory bulbs and brain in 1% CHAPS lysis buffer. Samples for western blot were prepared by adding 3x Loading Buffer (Life Technologies) containing 5 mM DTT and denatured for 5 minutes at 95°C. 30μg of proteins were loaded in to precast gels (Bio-Rad) along with a multicolour spectra high range protein ladder (Thermo Scientific). Gels were placed in a gel tank and submerged in 1x Running Buffer. Samples were electrophoresed for between 1-3 hours, depending on size of protein of interest at 120 volts. Immobilin-P membrane (Millipore) was activated with methanol (Sigma) followed by wet transfer of proteins in 1x ice cold Transfer Buffer. Transfer was carried out for 1 hour at 100 volts. Membranes were then blocked using 5% Marvel dry milk in 1x PBS. Antibody incubation was performed overnight at 4^o^C. The membrane was then washed and incubated with HRP tagged secondary antibody in PBS containing 2.5% dry milk in 1x PBS, with agitation at room temperature. Membranes were then washed and the proteins were visualised using Super Signal West Dura Extended Duration Substrate (Thermo Scientific) on light sensitive film (GE Healthcare) and developed on a Konica Minolta (SRX-101A) medical film developer.

### Plasmids, cloning and primers

The following plasmids were obtained from Addgene (Cambridge, MA): Tmed10, Gprin1 and Lat1. In order to perform *in vitro* validation of Na_V_1.7 interaction with Synaptotagmin-2, we first cloned the human gene for insertion into a mammalian expression plasmid. Synaptotagmin-2 was cloned from human dorsal root ganglion neuronal tissue. Whole mRNA was first reverse transcribed into a cDNA library for PCR. Following insertion into a TOPO vector, this was then used as the template to create the Synaptotagmin-2 insert, which was then successfully cloned into a pcDNA3.1 IRES-AcGFP plasmid using the Gibson assembly method. The Collapsin Response Mediator Protein 2 (Crmp2) gene was cloned from human dorsal root ganglia neuron mRNA and cloned into a pcDNA3.1 plasmid using Gibson assembly. The Scn3b mammalian expressing vector was used to investigate loss-of-function of Na_V_1.7 (Cox et al., 2006).

### Co-immunoprecipitation (Co-IP)

Mouse tissues used in immunoprecipitation experiments were chosen on the basis of known Na_V_1.7 expression, these include the following: Olfactory bulb, hypothalamus, spinal cord, dorsal root ganglia and sciatic nerve. All tissues from individual mice were either pooled or treated as individual tissue samples depending on experimental requirements. All tissues were flash frozen in dry ice immediately following dissection and stored at −80°C to avoid protein degradation. HEK293 cells stably expressing TAP-tagged Na_V_1.7 were harvested by trypsinisation followed by centrifugation at 800 rpm for 5 minutes. Cell pellets were stored at −80°C to avoid protein degradation. Samples were lysed in a 1% CHAPS lysis buffer containing a protease inhibitor cocktail and homogenised using ceramic zirconium oxide mix beads of 1.4 mm and 2.8 mm lysing kit and homogeniser (Precellys). A total of 10-25mg of protein was incubated with M2 Magnetic FLAG coupled beads (Sigma-Aldrich) for 2h at 4°C. The bead-protein complex was then washed with three cycles of 5 resin volumes of 1% CHAPS buffer and once with TEV-protease buffer (Invitrogen). The tagged protein was cleaved from the beads by the addition of TEV protease enzyme (Invitrogen) and incubated for 3 hours at 37°C to elute the protein complex. Sample eluate was then separated from the beads and stored at −80°C.

### Liquid chromatography-tandem mass spectrometry (LC-MS/MS) analysis

Proteins cleaved from Ni-NTA beads after affinity purification were tryptic digested following a FASP protocol (Wisniewski et al., 2009). In brief, proteins were loaded to 30 KDa filters (Millipore), then filter units were centrifuged at 14,000g for 15 min to remove other detergents. Two hundred μl of urea buffer (10 mM dithiolthreitol 8M urea (Sigma) in 0.1 MTris/HCl pH 8.5) were added to the filters and left at room temperature for 1 hour to reduce proteins. The filters were centrifuged to remove dithiolthreitol. Two hundred μl of 50 mM iodoacetamide in urea buffer were added to filters and left 30 min in the dark. The filters were centrifuged as before to remove IAA. Then the samples were buffer exchanged twice using 200 μl of urea buffer, and one more time using 200 μl of 50 mM NH4HCO3 in water. Forty μl of 50 ng/μl trypsin in 50 mM NH4HCO3 were added to filter, filters were vortexed briefly and proteins were digested at 37 °C for overnight. After tryptic digestion, the filters were transferred to new collection tubes, and the peptides collected by placing the filter upside down and spinning. The samples were acidified with CF3COOH and desalted with C18 cartridge (Waters). The pure peptides were dried by Spedvac (Millipore) and resuspended with 20 μl of 2% ACN, 0.1% FA. Five μl of samples were injected into Orbitrap velos mass spectrometry (Thermo) coupled to a UPLC (Waters) (Thézénas et al., 2013).

LC-MS/MS analysis was carried out by nano-ultra performance liquid chromatography tandem MS (nano-UPLC-MS/MS) using a 75 μm-inner diameter x 25 cm C18 nanoAcquity UPLC^TM^ column (1.7-μm particle size, Waters) with a 180 min gradient of 3 - 40% solvent B (solvent A: 99.9% H2O, 0.1% formic acid; solvent B: 99.9% ACN, 0.1% Formic acid). The Waters nanoAcquity UPLC system (final flow rate, 250 nl/min) was coupled to a LTQ Orbitrao Velos (Thermo Scientific, USA) run in positive ion mode. The MS survey scan was performed in the FT cell recoding a window between 300 and 2000 m/z. the resolution was set to 30,000. Maximum of 20 MS/MS scans were triggered per MS scan. The lock mass option was enabled and Polysiloxane (m/z 371.10124) was used for internal recalibration of the mass spectra. CID was done with a target value of 30,000 in the linear ion trap. The samples were measured with the MS setting charge state rejection enabled and only more than 1 charges procures ions selected for fragmentation. All raw MS data were processed to generate MGF files (200 most intense peaks) using the Proteowizard v.2.1.2476 software. The identification of proteins was performed using MGF files with the central proteomics facilities pipeline. Mus musculus (Mouse) database containing entries from UniProtKB was used in CPF Proteomics pipeline for data analysis. This pipeline combines database search results from three search engines (Mascot, OMSSA and X!tandem k-score). The search was carried out using the following parameters. Trypsin was the enzyme used for the digestion of the proteins and only one missed cleavage was allowed. The accepted tolerance for the precursor was 20 ppm and 0.5 Da for the fragment. The search encompassed 1+, 2+ and 3+ charge state, fixed modification for cysteine carbamidomethyl and variable modification for asparagine and glutamine deamidation, and methionine oxidation. All trypsin fragments were added to an exclusion list. False discovery rate was calculated by peptide/proteinprophet or estimated empirically from decoy hits, identified proteins were filtered to an estimated 1% FDR. The label-free analysis was carried out using the normalized spectral index (SINQ) (Trudgian et al., 2011). The mass spectrometry proteomics data have been deposited to the ProteomeXchange Consortium (http://www.proteomexchange.org/) via the PRIDE (Vizcaíno et al., 2016) partner repository with the dataset identifier PXD004926.

### Electrophysiology and patch clamp recordings

Whole cell patch clamp recordings were conducted at room temperature (21°C) using an AxoPatch 200B amplifier and a Digidata 1322A digitizer (Axon Instruments), controlled by Clampex software (version 10, Molecular Devices). Filamented borosilicate microelectrodes (GC150TF-10, Harvard Apparatus) were pulled on a Model P-97 Flaming/Brown micropipette puller (Sutter Instruments) and fire polished to a resistance of between 2.5-4 MOhm. Standard pipette intracellular solution contained: 10 mM NaCl, 140 mM CsF, 1.1 mM EGTA, 1 mM MgCl2, and 10 mM HEPES. The standard bathing extracellular solution contained: 140 mM NaCl, 1 mM MgCl2, 3 mM KCl, 1 mM CaCl2 and 10 mM HEPES. Both intracellular and extracellular solutions were adjusted to a physiological pH of 7.3. The amplifier’s offset potential was zeroed when the electrode was placed in the solution. After a gigaseal was obtained, short suction was used to establish whole-cell recording configuration. Errors in series resistance were compensated by 70 to 75%. Cells were briefly washed with extracellular solution before a final 2 mL of solution was transferred to the dish. Cells were held at −100 mV for 2 minutes before experimental protocols were initiated. Currents were elicited by 50 ms depolarisation steps from −80mV to + 80mV in 5mV increments. Compounds were added and mixed at the desired concentrations in extracellular solution before being added to the bath. Following addition of the compound protocols were repeated on previously unrecorded cells. All currents were leak substracted using a p/4 protocol. The following compounds were used in electrophysiology experiments: Lacosamide ((R)-2-acetamido-N-benzyl-3- methoxypropionamide) was obtained from Toronto Research Chemicals Inc (L098500) and Tetrodotoxin was obtained from Sigma-Aldrich (T8024). Incubation with Lacosamide was done for 5 hours prior to recording.

Voltage-clamp experiments were analysed using cCLAMP software and Origin (OriginLab Corp., Northampton, MA) software programs. Current density-voltage (pA/pF) analysis by measuring peak currents at different applied voltage steps and normalised to cell capacitance. Voltage dependent activation data was fitted to a Boltzmann equation y = (A2 + (A1 – A2)/(1 + exp((Vh – x)/k))) * (x – Vrev), where A1 is the maximal amplitude, Vh is the potential of half-maximal activation, x is the clamped membrane potential, Vrev is the reversal potential, and k is a constant. All Boltzmann equations were fitted using ORIGIN software.

### Na_V_1.7 interaction protein selection and function analysis

Candidate proteins that may interact with Na_V_1.7 were selected by two criteria: a) Present in at least two knock-in biological experiments but absent from wild type experiments. b) Present in more than three knock-in and more than one wild type experiments, the ratio of average abundance is more than 1.5 fold increased in knock-in experiments as compared with wild type experiments. Further cellular component and function classification were performed on PANTHER Classification System (11.0). Ingenuity pathway analysis (IPA) (QIAGEN) was used to elucidate pathways and protein interaction networks using candidate proteins.

### Southern blot analysis

The genomic DNA was extracted from either ES cells or tails of mice following the procedures as described (Sambrook and Russell, 2001). The probes for Southern blot were amplified by PCR using mouse genomic DNA isolated from C57BL/6 as a template and purified with a Qiagen Gel Purification Kit. The restriction enzymes *Stu*I*, BspH*I and *Psi*I were used to digest genomic DNA for either wild-type and knock-in bands. The sizes of wild-type and knock-in bands is shown in Fig 2B. The primers used to create probes (5’ external probe: 768 bp; 3’ external probe: 629 bp) included:

5’PF (5’probe, forward) - ACCAAGCTTTTGATATCACCATCAT
5’PR (5’probe, reverse) - CAACTCGAGAACAGGTAAGACATGACAGTG
3’PF (3’probe, forward) - TTTAAGCTTCCTGCCCCTATTCCTGCT
3’PR (5’probe, reverse) – TTAGGATCCATGCACTACTGACTTGCTTATAGGT

### Statistical Analysis

Statistical analysis was performed using either repeated-measures ANOVA with Bonferroni post hoc testing or Unpaired Student’s *t*-test as described in the results or figure legends. The GraphPad Prism 6.0 was used to perform the statistical analysis. All data are presented as mean ± SEM and significance was determined at *p* < 0.05.

## Acknowledgments

We are grateful to Professor Seth GN Grant and Dr. Esperanza Fernandez for the TAP-tag plasmids. We thank the Mass Spectrometry Laboratory of Target Discovery Institute at University of Oxford for performing the Mass spectrometry analysis. We thank Dr. Massimo Signore who performed electroporation and blastocysts microinjection of ES cells in JP Martinez-Barbera’s research group, ICH, UCL. This study was supported by the Wellcome Collaborative Award 200183/Z/15/Z (to J.W., J.Z., J.C. and S.G.) and Investigator Award 101054/Z/13/Z (to J.W.), the Medical Research Council (MRC) Grant G091905 (to J.W.) and MRC CDA G1100340 (to J.C.), the Deutsche Forschungsgemeinschaft Grants SFB 894/A17 (to F.Z.) and PY90/1-1 (to M.P.)

